# Photoswitchable endocytosis of biomolecular condensates in giant vesicles

**DOI:** 10.1101/2024.01.10.574984

**Authors:** Agustín Mangiarotti, Mina Aleksanyan, Macarena Siri, Reinhard Lipowsky, Rumiana Dimova

## Abstract

Interactions between membranes and biomolecular condensates can give rise to complex phenomena such as wetting transitions, mutual remodeling, and endocytosis. In this study, we demonstrate a light-triggered manipulation of condensate engulfment using giant vesicles containing photoswitchable lipids. UV irradiation increases the membrane area, facilitating a rapid condensate endocytosis, which can be reverted by blue light. The affinity of the protein-rich condensates to the membrane and the reversibility of the engulfment processes is quantified from confocal microscopy images. The degree of engulfment, whether partial or complete, depends on the initial membrane excess area and the relative sizes of vesicles and condensates. Theoretical estimates suggest that utilizing the light-induced excess area to increase the vesicles-condensate adhesion interface is energetically more favorable than the energy gain from folding the membrane into invaginations and tubes. Our overall findings demonstrate that membrane-condensate interactions can be easily and quickly modulated via light, providing a versatile system for building platforms to control cellular events and design intelligent drug delivery systems for cell repair.

## 1. Introduction

Biomolecular condensates are specialized, membraneless cellular compartments formed through the dynamic and reversible assembly of proteins, nucleic acids and other biomacromolecules within the cytoplasm or nucleus of a cell [1]. They are involved in a wide range of cellular processes including gene transcription and translation [2], signal transduction [3], stress response [4], protein quality control [5], cell division [6]. Lately, condensate-membrane interactions were shown to be crucial for the formation of tight junctions [7], transport of stress granules [8], signal transduction in T-cells [3], and endocytosis [9]. The associated morphologies are governed by membrane wetting transitions [10] that can be tuned via changes in the salinity of the milieu or the membrane composition [10b, 11]. *In vitro* studies have demonstrated that condensate endocytosis can occur when sufficient membrane area is available [10b, 11]. Furthermore, coacervates of various compositions can be completely engulfed by lipid vesicles via altering the membrane charge and increasing the strength of droplet-membrane interaction [11]. Control over these processes allows the development of synthetic cells [12], which can be used for therapeutic purposes in biotechnology by designing effective drug delivery platforms [13].

Introducing biocompatible molecular photoswitches into bio-systems provides an additional leverage for fast and reversible manipulation of cellular processes through light [14]. Here, we construct a photoswitchable biomimetic system for condensate endocytosis using giant unilamellar vesicles (GUVs) and glycinin protein condensates. Giant vesicles are cell-sized, biomembrane compartments with increasingly broad spectrum of applications [15], and glycinin is a well-studied storage protein from the soybean that constitutes a robust model for protein condensation [16]. UV exposure of GUVs composed of 1-palmitoyl-2-oleoyl-glycero-3-phosphocholine (POPC) and a photoswitchable azobenzene phospholipid analog (azo-PC) can trigger reversible increase in membrane area upon trans-to-cis isomerization [17]. This suggests that light can finely manipulate the properties of photolipid-doped cell models and liposomal drug carriers, offering potential therapeutic applications for addressing cellular disorders. Employing GUVs with photoswitches we demonstrate light-triggered endocytosis of glycinin condensates in an instantaneous and reversible manner.

## 2. Results and Discussion

### 2.1. Reversible partial and complete engulfment of protein condensates under the influence of light

When the photoswitchable azo-PC lipids are exposed to UV-light (365 nm), *trans*-to-*cis* photo-isomerization occurs (Figure 1a), which effectively increases the area per molecule and the total vesicle area accordingly. Photoisomerization in membranes containing 50 mol% azo-PC results in a substantial area increase of ∼18 % [17]. We prepared POPC:azo-PC (1:1 molar ratio) GUVs labeled with 0.1 mol% ATTO-647N-DOPE and observed them with confocal microscopy. Under UV light, the GUVs grow in size and the generated excess membrane area quickly transforms into internal nanotubes (Figure 1b,c). This process can be reversed by blue irradiation (450 nm), see Movie S1. The formation of nanotubes is caused by the buffer asymmetry across the GUV membrane (sucrose solution inside and isotonic sodium chloride solution outside) resulting in high negative spontaneous curvature stabilizing the tubes [18]. The sodium chloride solution was required for condensate formation (see SI for details) while the internal sucrose solution osmotically stabilizes the GUV ensuring volume conservation.

**Figure 1.**
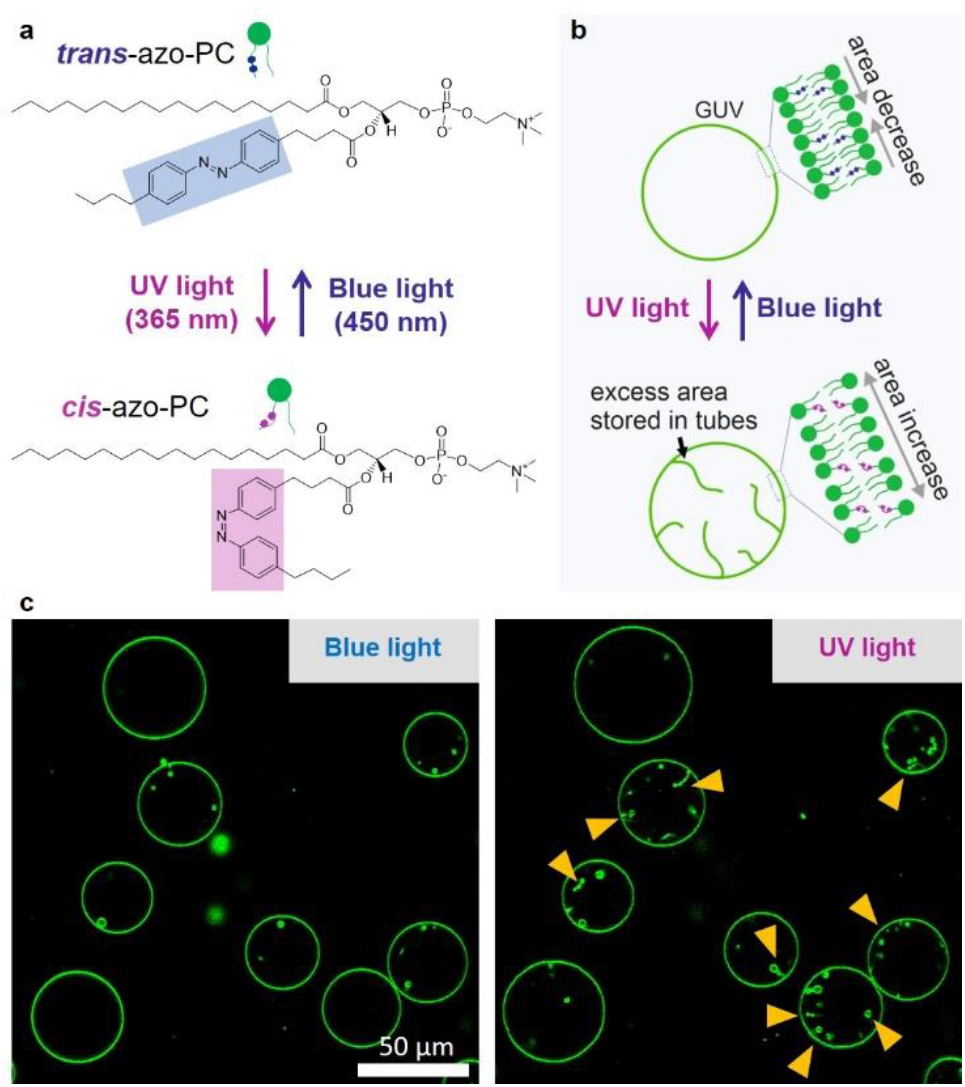
Irradiation of photoswitchable azo-PC causes reversible area increase and morphological changes in GUVs. (a) Molecular structures of *cis* and *trans* azo-PC photoswitchable isomers. (b) Schematic illustration of the reversible membrane area changes in GUVs under UV and blue light. The excess area generated under UV irradiation can be stored in nanotubes. (c) Confocal cross-sections showing membrane area change of POPC:azo-PC (1:1) GUVs labelled with 0.1 mol% Atto-647N-DOPE upon *trans*-*cis* photoisomerization and extensive nanotube formation in the GUV lumen (arrowheads); see Movie S1.

To probe whether the condensate-membrane interaction can be tuned by light, we placed the vesicles in contact with glycinin condensates formed at 150 mM NaCl and labelled with the water-soluble dye Sulforhodamine B (SRB). Upon adhesion, the condensates deform the GUV membrane (Figure 2), as previously shown [10b]. Under UV irradiation, the *trans-*to*-cis* photoisomerization of azo-PC results in fast membrane expansion accompanied by increasing adhesion zone. We observed two outcomes depending on the relative sizes of the interacting vesicle and condensate (Figure 2). For larger condensates, with radius (*R*_*cond*_) comparable to or exceeding the vesicle radius (*R*_*GUV*_), the generated excess membrane is consumed to partially engulf the droplet, see Figure 2a,b and Movie S2. For smaller condensates (*R*_*GUV*_>>*R*_*cond*_), the UV-induced excess area of the GUV allows complete engulfment, i.e. endocytosis of the condensate (Figure 2c,d and Movie S3). Images of the membrane channel and the 3D projection show that the condensate is fully enclosed by the membrane, confirming endocytosis (Figure S1). Note that to allow reversibility of endocytosis, the membrane adhering to the droplet must be connected to the mother vesicle by a closed (nanometric) membrane neck [19]. This membrane neck can be subsequently cleaved via scaffolding proteins [20] or low density of membrane bound-proteins inducing large spontaneous curvature that generates a constriction force [21].

**Figure 2.**
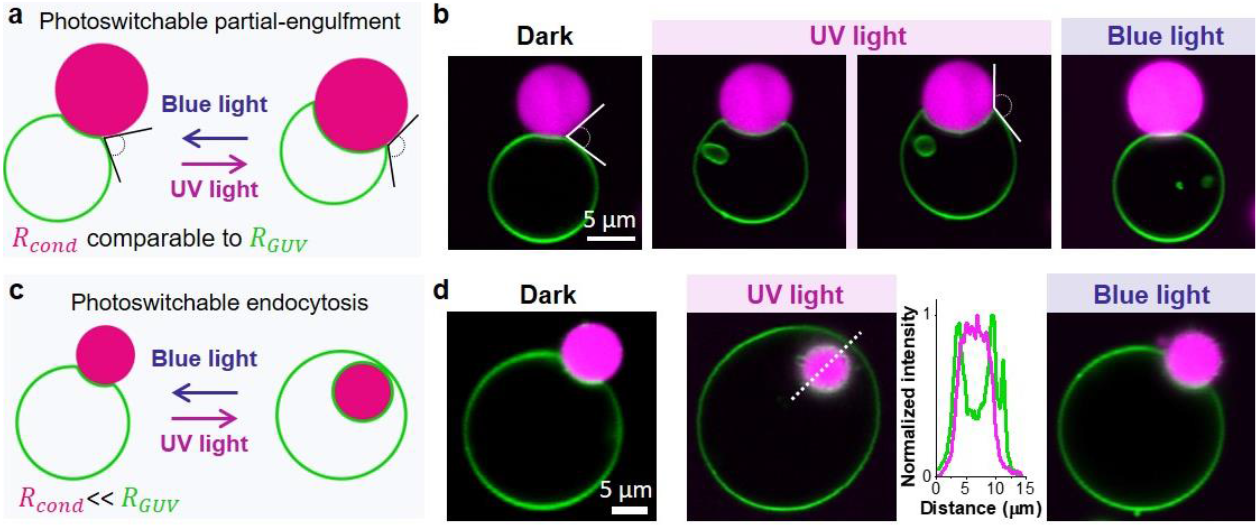
Light-induced engulfment of condensates. (a,b) Condensates with sizes comparable to that of the GUV experience reversible partial engulfment (schematic and confocal cross-sections) mediated by UV/blue irradiation, see Movie S2 for time-lapse recording. The contact angle between the condensate and the membrane changes upon photoisomerization as highlighted. (c,d) For smaller condensates, photoisomerization leading to sufficient excess area leads to complete engulfment, which is reversible (Movie S3). The intensity profile corresponds to the white dashed line shown in the second image in (d) showing the position of the condensate (magenta) wrapped by the membrane (green).

In addition, a single GUV can engulf multiple condensates, as shown in Figure S2a. However, this case requires more available membrane area, and the adhesion of multiple condensates might increase the membrane tension [11, 22], reducing the likelihood of the engulfment of many droplets.

As shown in Figure 2, both, the complete and partial engulfment processes can be quickly reversed by blue light exposure, see Figures S2-S4 and Movies S4-S8 for more examples including large-field images. We emphasize that the protein structure remains unaltered after UV irradiation as demonstrated by Fourier-transform infrared spectroscopy (see Figure S5, Table S1 and Experimental section), indicating that the observed changes in membrane-condensate affinity are solely due to the light-induced changes in the membrane.

### 2.2. Quantifying the condensate affinity to the membrane from the system geometry

In the case of the partial engulfment of condensates by GUVs, the contact angle between the membrane and the condensate (Figure 2a,b) changes, suggesting altered affinity. Note that measuring only this apparent contact angle may provide an inaccurate understanding of the interaction. Assessing it solely from confocal cross-sections where either the condensate center or the vesicle centers are not in the plane of the image (out of focus) results in an incorrect system geometry description. To obtain accurate information, it is crucial that the rotational axis of symmetry of the vesicle-droplet system lies in the image plane of the projected image. This often necessitates the acquisition and reorientation of a 3D image of the vesicle and droplet., see e.g. [10b].

The changes in the affinity of the condensate to the membrane can be precisely quantified by measuring the geometric factor Φ = (sin θ_*e*_ − sin θ_c_)/sin θ_*i*_, which is obtained from the microscopically measured contact angles facing the external phase, θ_*e*_, the vesicle lumen, θ_*i*_, and the condensate interior, θ_c_. Note that while the membrane shape appears to have a kink at the three-phase contact line, the detailed structure of this kink corresponds to a highly curved membrane segment that is resolvable only with super-resolution microscopy [23].

The geometric factor is a dimensionless quantity reflecting the tensions in the system, namely the interfacial tension Σ_ce_ and the two mechanical tensions of the membrane segments in contact with the external phase, 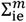, and with the condensate, 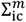 (see details in Figure 3a). It can adopt values between -1, corresponding to complete wetting and spreading of the condensate on the membrane surface, and 1 corresponding to no wetting where both vesicles and condensate remain spherical. The intermediate values, −1 < Φ < 1, reflect the case of partial wetting. While the contact angles reflect the specific geometry of a vesicle-condensate couple and can exhibit broad variations in the sample, the geometric factor is a material property of the condensate-membrane system and is constant over the whole population. It reflects the wetting affinity and is independent of condensate-vesicle geometry as well as droplet and vesicle sizes [10b].

**Figure 3.**
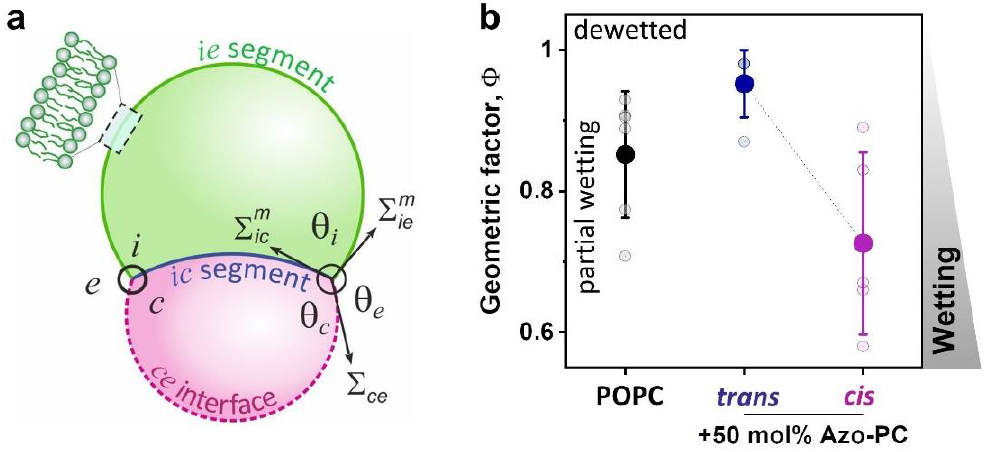
System geometry and contact angles used to determine the geometric factor, which shows increased wetting affinity upon *trans*-to-*cis* isomerization. (a) For partial wetting morphologies of condensate-membrane systems, the contact interface between the condensate (magenta) and the membrane (green) partitions the membrane into the *ie* and *ic* segments, with the contact angles θ_i_ + θ_e_ + θ_c_=360°. The interfacial tension Σ_ce_ and the mechanical tensions 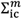 and 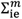 within the two membrane segments are balanced, see [10b, 22]. (b) Experimental data for the geometric factor Φ = (sin θ_*e*_ − sin θ_c_)/sin θ_*i*_ for pure POPC GUVs or POPC:azo-PC 1:1 GUVs in contact with glycinin condensates at 150 mM NaCl. The geometric factor reaches the maximum value of Φ = +1 for complete dewetting (no interaction between the condensate and the vesicle) and partial wetting for lower values. The drop in Φ for the *trans-*to*-cis* isomerization of the azo-PC lipids indicates that the affinity between the condensates and the membrane is increasing.

Figure 3b shows that the geometric factor for pure POPC and POPC:azo-PC membranes in the *trans* state are similar, but *trans-*to*-cis* photo-isomerization alters it. The data indicate that the condensate-membrane affinity increases upon *trans-*to*-cis* photo-isomerization of azo-PC.

### 2.3. Light-triggered condensate engulfment and reversibility kinetics

Next, we assessed the light-triggered endocytosis and partial engulfment over multiple cycles of UV and blue light irradiation. To quantify the light-induced engulfment and release of the protein condensates from GUVs containing 50 mol % azo-PC, the degree of the penetration of the protein condensates in the vesicles (penetration depth, *p*) was calculated. For this, we used confocal cross-sections at different levels perpendicular to the optical axis (z-axis) within the sample exposed to UV and blue illumination for two photo-switching cycles. The definition of *p* is based on the study of Dietrich et al. [24], which addresses the adhesion dynamics of spherical solid particles to lipid vesicles. The penetration distance of the protein condensate, denoted as *d*, is measured along the axis of rotation. It represent the distance from the estimated outer rim of the interpenetrated lipid vesicle to the interface between the protein condensate and the vesicle membrane, as illustrated in Figure 4a; see also the Experimental section.

**Figure 4.**
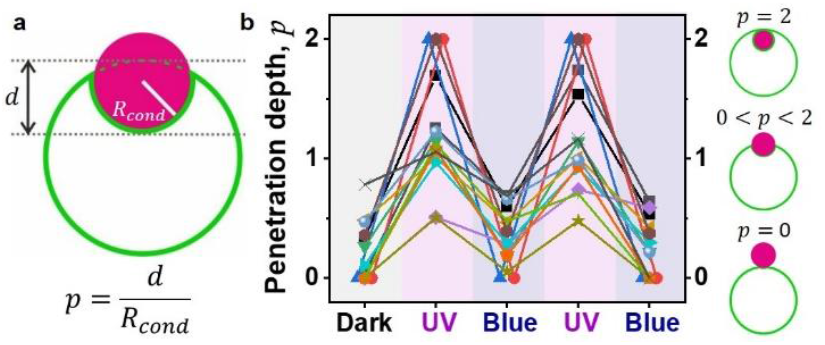
Reversible endocytosis. (a) Sketch showing the measured parameters for calculating the rescaled penetration depth, *p*. (b) The penetration depth of condensates into vesicles, *p*, is fully reversible for multiple photoswitching cycles (up to six cycles were tested). Each symbol represents a single vesicle-condensate pair.

Figure 4b shows data for the penetration depth where *p* = 2 corresponds to endocytosis and 0 < *p* < 2 reflects partial engulfment. The value of *p* depends on the initial excess area and relative condensate-vesicle sizes. It alternates with photoswitching cycles of UV and blue light and is fully reversible. Both light-triggered partial and complete engulfment processes are characterized by fast kinetics in the milliseconds (*cis*-to-*trans*) to a few seconds range (*trans*- to-*cis*), see Figure 5. This is consistent with the notion that the *trans* state is more stable. These light-triggered kinetics are faster than passive engulfment, which can take from seconds to minutes, or even hours depending on the particular condensate-vesicle system [11].

**Figure 5.**
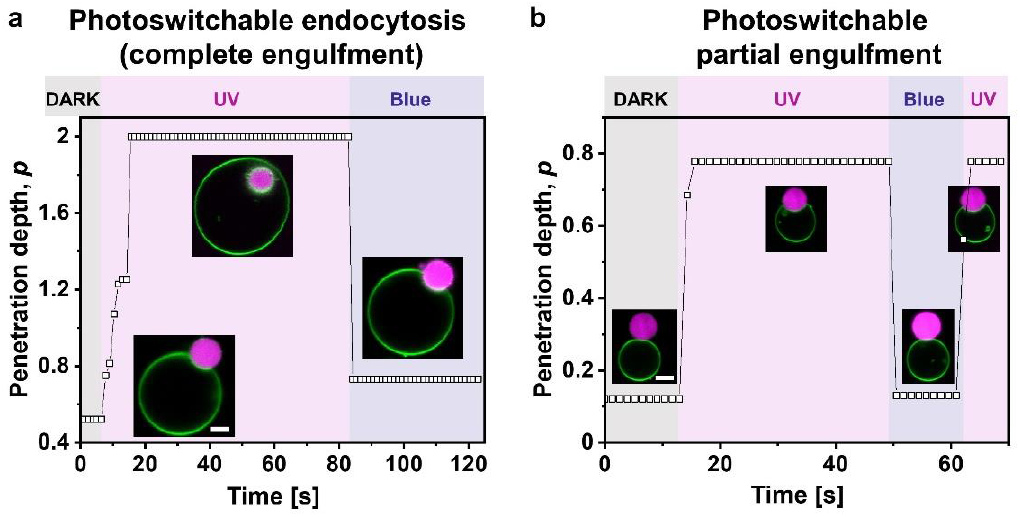
Penetration depth, *p* vs time showing the kinetics of photoswitchable endocytosis (a) and partial-engulfment (b). Scale bars are 5 µm. The *trans*-to-*cis* photoisomerization effect is completed in 2-9 seconds, while the *cis*-to-*trans* photoisomerization occurs at times faster than the frame rate of the image acquisition (below 650 milliseconds).

### 2.4. Recruiting excess membrane area for condensate adhesion is energetically more favorable than membrane tubulation

As shown in Figure 1, the excess vesicle area generated by the *trans-*to*-cis* photoisomerization of azo-PC can be stored in tubes. When in contact with a condensate, this excess area goes to the membrane-condensate interface enabling partial or complete endocytosis of the condensate (Figures 2, 5). However, we observed that nanotube formation can also take place during and after the engulfment of condensates. We questioned whether all area generated by *trans-*to*-cis* photoisomerization is transferred to the condensate interface. This required precise determination of the UV-induced area increase, which we assessed from electrodeformation of GUVs [25] in the absence of condensates. The vesicles were exposed to an alternating current (AC) field to pull out membrane fluctuations and inducing elliptical deformation [26]. This allows the precise measure of the total vesicle area from its geometry, see the Experimental section. Subtracting this initial membrane area from the membrane area under the influence of both UV light and AC field, yields UV-induced area increase of 18±2%, (see Figures S6-S8), which is in good agreement with reported data for slightly different solution conditions [17]. We then compared this expected absolute area increase for individual vesicles wetted by a condensate with the change in apparent area of the vesicles measured directly from the microscopy images as the area sum of the bare vesicle membrane segment and the membrane segment in contact with the condensate (i.e. excluding the area of possible tubular structures), see Figure 6. Provided all UV-induced excess area is consumed to expand the membrane-condensate interface, the data should fall on a line described by *y* = *x*. However, many datapoints are distributed above this line, indicating that the excess area does not only accumulate at the membrane-condensate interface, but is also being stored in membrane nanotubes.

**Figure 6.**
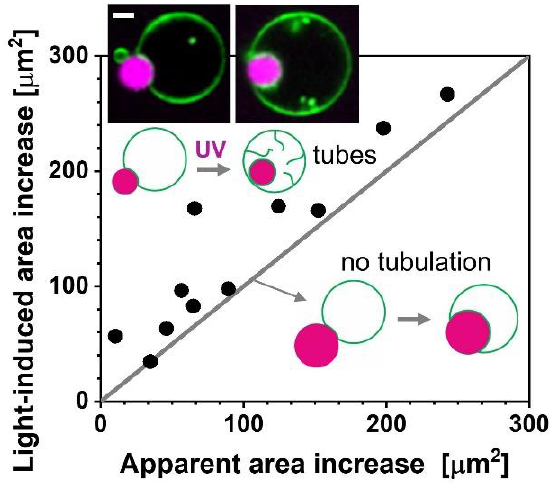
Light-induced area increase as expected from electrodeformation experiments (Figures S9, S10) versus apparent area increase (area of the spherical segments of the bare and wetted membrane excluding tubes) for POPC:Azo-PC 1:1 GUVs in contact with condensate droplets. Data falls at the *y* = *x* gray line when all the excess area is transferred to the membrane-condensate contact area, as shown in the lower sketches. Data lying above this line corresponds to vesicles in which part of the excess area is stored in nanotubes (upper sketch and example images; scale bar 5 µm).

High salt and sugar concentrations are known to modify the membrane structure and properties [27]. Under high salt/sugar asymmetry across the vesicle membrane, ion adsorption to the outer leaflet produce negative membrane spontaneous curvature, which stabilizes inward tubes [18]. Note that in the absence of this salt asymmetry, UV irradiation only produces large membrane fluctuations in the GUVs but no tube formation, see Figure S10. Thus, the total area increase due to UV irradiation of the vesicles (as plotted in Figure 6) represents the sum of the outer spherical vesicle membrane and the membrane area stored in tubes.

We also compared this area change to the change in the area of the membrane segment wetted by the condensate before and after UV illumination (Figure S9). Similar trend was observed. The observations imply that nanotube formation competes with transferring the excess area to the membrane-condensate interface, and that the final morphology could depend on the initial geometry of the vesicle-condensate pair (note that the preparation method leads to vesicles with different area-to-volume ratios).

Next, we theoretically estimated the energetic gain of transferring the excess area to the vesicle-condensate interface and compared it to that arising from the formation of nanotubes. To estimate the energy gain arising from tubulation, we consider a cylindrical tube of area Δ*A* stabilized by membrane spontaneous curvature *m*. The tube is characterized by radius 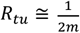. The gain of bending energy, Δ*E*_*be*_ associated with the transfer of the membrane area Δ*A* from the weakly curved mother vesicle to the nanotube is given by

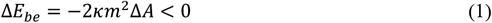

where *κ* is the membrane bending rigidity. For membranes asymmetrically exposed to sugar and sodium chloride solutions as studied here, *m* ≈ 9 µm^−1^ as measured in Ref. [18]. The bending rigidity for POPC membranes containing 50 mol% of azo-PC is *κ* ≈ 10 *k*_B_*T* [17], where *k*_B_*T =* 4.1x10^-21^ J is the thermal energy. The excess area Δ*A* available for tube formation is simply the difference between the light-induced area change *A*_*li*_ and the apparent area *A*_*app*_ plotted in Figure 6, where we see that Δ*A* = *A*_*li*_ − *A*_*app*_ ranges between 0 and roughly 160 µm^2^. Taking 80 µm^2^ as a mean value of Δ *A*, we obtain for the bending energy gain Δ*E*_*be*_ ≈ −1.3×10^5^ *k*_B_*T*.

As long as the *ce* interface of the droplet (see Figure 3a) is not completely covered by the vesicle membrane, the photo-induced excess area Δ*A* can be alternatively used to increase the contact area between droplet and membrane. The adhesion energy per unit area is given by

ΦΣ_*ce*_Δ*A* with Φ = 0.75 when the membrane is exposed to UV light and the azo-PC lipids attain their *cis*-conformation (Figure 3b). As we cover the area Δ*A* of the droplet surface, we reduce its interfacial free energy by Σ_*ce*_Δ*A*. Therefore, the gain in adhesion energy is equal to

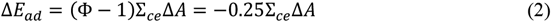

where the numerical value Φ = 0.75 for *cis*-azo-PC has been used in the second equality. The interfacial tension has the value Σ_*ce*_ ≈ 0.5 mN/m [10b] which implies Δ*E*_*ad*_ = − 2.4×10^6^ *k*_B_*T* for Δ*A =* 80 µm^2^. Comparing the gain Δ*E*_*ad*_ in adhesion energy with the gain Δ*E*_*ad*_ ≈ −1.3×10^5^ k_B_T in bending energy as caused by tubulation, we conclude that the gain in adhesion energy exceeds the gain in bending energy by more than one order of magnitude. As a consequence, the vesicle membrane will continue to spread over the droplet until this droplet is completely engulfed. Any additional excess area created by the light-induced isomerization will be stored in membrane nanotubes. Indeed, subsequent inspection of the vesicle-condensate images demonstrated that data points in Figure 6 located above the line with slope 1 correspond to vesicles where the condensates are completely engulfed and the remaining excess area engages in the formation of nanotubes (see confocal images in the inset).

## 3. Conclusions

In summary, our work shows that light can be used as a facile, inexpensive, and sustainable tool for efficiently tuning the membrane-condensate interactions in fast and reversible manner. By using GUVs containing the azo-PC photolipid as minimalistic artificial cells, we effectively generated and characterized light-induced membrane-condensate wetting transitions leading to fast reversible endocytosis (within a few seconds) over multiple photoswitching cycles. By combining theoretical studies with experimental observations, we have elucidated the interaction mechanisms between protein condensates and lipid membranes leading to engulfment and membrane morophology changes. The application of these results could be extended to different condensate systems and membrane compositions, provided that there is partial wetting between the condensate and the vesicles. The photoswitchable system presented here provides a promising platform for the development of synthetic cells, and versatile drug delivery systems with applications in photo-pharmacology.

## 4. Experimental Section

### Materials

The phospholipids 1-stearoyl-2-[(E)-4-(4-((4-butylphenyl)diazenyl)phenyl)butanoyl]-sn-glycero-3-phosphocholine (azo-PC) and 1-palmitoyl-2-oleoyl-sn-glycero-3-phosphocholine (POPC) were purchased from Avanti Polar Lipids, Alabaster, AL, USA. The lipid 1,2-dioleoyl-sn-glycero-3-phosphoethanolamine labeled with Atto 647N (Atto-647N-DOPE) was obtained from ATTO-TEC GmbH, Siegen, Germany. Sucrose, glucose and sodium chloride (NaCl) were purchased from Merck, Germany. The fluorescent dye Sulforhodamine B (SRB) was obtained from ThermoFisher Scientific, Massachusetts, USA. Bovine serum albumin (BSA) was purchased from Merck, Germany. Stock solutions of the phospholipids and the dye-conjugated lipid were prepared in chloroform solution to concentration of 4 mM and stored at -20°C until usage. Indium-tin oxide (ITO)-coated glass plates were purchased from PGO GmbH, Iserlohn, Germany. An Agilent 33220A 20 MHz Function/Arbitrary Waveform Generator from Agilent Technologies, USA was used for GUVs electroformation. The microscopic observations were done either with commercially available Eppendorf electrofusion chambers (Germany) or home-made chambers assembled from 22×40 mm^2^ and 22×22 mm^2^ cover slides purchased from Knittel Glass (Germany). Cover slides were rinsed with ethanol and distilled water and then passivated with a 2 mg/mL BSA solution. An Osmomat 3000 osmometer (Gonotec GmbH, Berlin, Germany) was used to measure solutions osmolarities.

All commercially available chemicals and solvents were used without further purification. In order to prevent any dust or dirt, all the glassware was rinsed with ethanol and chloroform, and then dried under inert atmosphere before usage.

### Vesicle preparation

Giant unilamellar vesicles (GUVs) were prepared at room temperature (23° C) by the electroformation method [15b]. An equimolar solution of azo-PC and POPC including 0.1 mol% Atto-647N-DOPE was prepared in chloroform to a final concentration of 4 mM. In order to create a thin lipid film, 14 µL of this lipid solution was first spread on a pair of electrically conductive, ITO-coated glass plates and then the majority of the chloroform was evaporated by exposing the plates to a stream of N_2_. For removal of solvent traces, the plates were also subsequently placed under vacuum for two hours. A chamber was assembled using a rectangular Teflon spacer of 2 mm thickness sandwiched between the ITO-glass plates. The chamber was filled with a solution of 300 mM (300 mOsmol/kg) sucrose to hydrate the lipid film. Electroswelling was induced by applying a sinusoidal alternating current (AC) electric field at 10 Hz frequency with a 1.6 V (peak to peak) amplitude for 1 hour in the dark. GUVs were then transferred to light-protective glass vials for storage at room temperature and used the same day.

For the GUV electrodeformation studies, GUVs were swelled in a 100 mM sucrose and 0.5 mM NaCl solution and then were 8-fold diluted in a 105 mM glucose solution. The presence of the small amount of salt ensures higher conductivity of the internal solution compared to the external one, and resulting in prolate deformation of the GUVs under AC field [26]. The control experiments of azo-PC GUVs in low sugar concentrations and in the absence of any salts in the external GUV medium were performed by harvesting GUVs in 100 mM sucrose solution and 1:1 dilution into 105 mM glucose solution for the confocal microscopy observations.

### Protein purification and condensate formation

Glycinin was purified as described by Chen et al [16a]. Briefly, defatted soy flour was dispersed 15-fold in water by weight and adjusted to pH 7.5 with 2 M NaOH. After centrifugation at 9000×g for 30 min at 4°C, dry sodium bisulfite was added to the supernatant (0.98 g/L). The pH of the solution was adjusted to 6.4 with 2 M HCl, and the obtained turbid dispersion was kept at 4 °C overnight. Next, the dispersion was centrifuged at 6500×g for 30 min at 4 °C. The glycinin-rich precipitate was dispersed 5-fold in water, and the pH was adjusted to 7. The glycinin solution was then dialyzed against Millipore water for two days at 4 °C and then freeze-dried to acquire the final product with a purity of 97.5% [16a].

To form the condensates, a 20 mg/mL glycinin solution at pH 7 was freshly prepared in ultrapure water and filtered with 0.45 µm filters to remove any insoluble materials. Then, the desired volume of the glycinin solution was mixed with the same volume of a 300 mM NaCl solution to achieve a solution with final concentrations of 10 mg/mL glycinin and 150 mM NaCl. The condensates were labelled by including 10 µM SRB dye prior to condensate formation.

### Condensate-vesicle suspensions

First, the vesicle suspension was diluted 1:10 in a 150 mM NaCl solution. Then, the condensate suspension was diluted 1:4 and added to the vesicle suspension at a 15% v/v (corresponding to a final protein concentration of 0.4 mg/mL). After gently mixing the vesicle-condensate suspension, an aliquot of 10 µL was placed on a coverslip for confocal microscopy and a chamber was formed using a round spacer and closed with a coverslip.

### Confocal microscopy imaging and irradiation conditions

UV-induced morphological changes of azo-PC GUVs as well as the engulfment of protein condensates into GUVs were monitored through a Leica TCS SP8 scanning confocal microscope (Wetzlar, Germany) using either a 40× (0.60 NA) air or 63× (1.2 NA) water immersion objectives. The pinhole size during the experiment was set to 1 AU (Airy unit) and the scanning speed was 400 Hz in bidirectional mode. Time-lapse imaging was performed at a frame rate of 650 ms/frame. SRB was excited with a 561 nm laser and the emission signal was collected with HyD (hybrid) detector in the 573-626 nm range. Atto-647N-DOPE was excited with a HeNe 633nm laser and the emission signal was collected with a HyD detector in the range 645-785 nm. In order to induce *trans-*to*-cis* photoisomerization of azo-PC in the GUVs, an external UV-LED lamp (365 nm wavelength; “UV light”) with a maximum power intensity of 20 mW cm^-2^ (Roschwege, Germany) was attached to the condenser of the confocal microscope. The reversed azo-PC photoisomerization (*cis-*to*-trans)* was generated by simultaneously using 458 nm and 476 nm lasers at 50% intensity (“blue light”).

### Phase contrast microscopy imaging and irradiation conditions

Electrodeformation of azo-PC GUVs was monitored under phase contrast mode of an inverted Axio Observer D1 microscope (Zeiss, Germany), equipped with a PH2 40x (NA 0.6) objective. Images were acquired with an ORCA R2 CCD camera (Hamamatsu, Japan). The GUVs were placed either in an Eppendorf electrofusion chamber or a home-made chamber with approximate thickness of 8 mm or 1 mm, respectively; see Figure S6. To induce UV irradiation, the light from the HBO 100W mercury lamp was used in epi-illumination mode and was collected through a 365 nm DAPI filter. For blue irradiation, the light from the mercury lamp was applied through a 470/40 nm filter. The irradiation power of the HBO lamp was measured with a LaserCheck power meter (Coherent, USA) at the position of the sample and recorded as 60 mW cm^-2^ for the UV filter and 26 mW cm^-2^ for the blue filter.

### Penetration depth analysis and irradiation

The penetration depth of the protein condensate into GUV, *p*, reflects the degree of insertion of the droplet inside the vesicle and is calculated as:

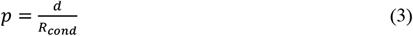

where *R*_*cond*_ and *d* are defined in Figure 4a. For each interacting condensate-GUV pair, *d* was measured from confocal screenshots using the Fiji software at 3 illumination conditions. In the initial illumination condition, “dark”, we used the 561 nm and 633 nm lasers to only excite the fluorescent dyes labeling the condensate and the membrane. In addition to these two lasers, an external UV LED at 365 nm was used to promote the *trans*-to-*cis* isomerization of azo-PC, and we refer to this condition as “UV-light”. Switching off the UV LED was followed by the immediate exposure of the sample to 458 nm and 476 nm lasers, which we refer to as “blue-light”. Then, the measured *d* and *R*_*cond*_ values were used in Equation 3 to calculate the *p* values at the three conditions (dark, UV, blue) for 2 photo-switching cycles. Origin Pro was used for plotting the penetration depth values from 15 protein condensates interacting with 8 GUVs (Figure 4b).

### Fourier-transform infrared (FTIR-ATR) spectroscopy

Spectra were recorded on an infrared microscope AIM-90000 (SHIMADZU, Germany) equipped with an ATR objective. First, a 3 µL aliquot of each sample was spread on glass slides, and dried until a film formed with N_2_. A second aliquot was spread on glass slides and irradiated for 10-20 seconds with the above-mentioned UV LED, before drying it with N_2_ until a film was formed. The ATR objective was placed pressuring the sample to acquire the protein spectra. Measurements consisted of an average of 64 scans recorded at 25°C with a nominal resolution of 4 cm^−1^. The spectra were processed using Kinetic software developed by Dr. Erik Goormaghtigh of the Structure and Function of Membrane Biology Laboratory (Université Libre de Bruxelles, Belgium). The spectra were analyzed in the amide I’ region of the protein (1700 and 1600cm^−1^). The spectra were deconvoluted using Lorentzian deconvolution factor with a full width at the half maximum (FWHM) of 30 cm^−1^ and a Gaussian apodization factor with a FWHM of 16.66 cm^−1^ to obtain a line narrowing factor K = 1.8. Band assignment was performed using the deconvoluted and second derivative spectra of each sample in the amide I’ region. These were the initial parameters for an iterative least square curve fit of the original IR band (K=1) using mixed Gaussian/Lorentzian bands. The bounds for the peak positions of each identified individual component were within ±2 cm^−1^ of the initial value. The FWHM input values are described in detail in Table S1. FTIR-ATR spectra and analysis of the secondary structure content of glycinin condensates before and after the UV illumination are demonstrated in Figure S5.

### Vesicle electrodeformation

Electrodeformation experiments to determine the membrane area changes associated with azo-PC isomerization were performed using both, a Leica TCS SP8 scanning confocal microscope (Wetzlar, Germany) equipped with a HC PL FLUOTAR L 40x (0.60 NA) objective, and an inverted microscope in phase contrast mode Axio Observer D1 (Zeiss, Germany) equipped with a PH2 40x (0.6 NA) objective and an ORCA R2 CCD camera (Hamamatsu, Japan). GUVs were observed in a commercial Eppendorf electrofusion chamber (Eppendorf, Germany) or a home-made chamber (see Figure S6) to compare the effect of chamber thickness on the penetration of UV-light through the sample and observation of light-induced changes on the vesicles in the sample. The Eppendorf chamber (see Figure S6a) contains two parallel cylindrical platinum electrodes 92 µm in radius, located 500 µm apart from each other. There the GUVs were exposed to an AC field (1 MHz, 5 V peak to peak amplitude), as previously described [26, 28]. The home-made chamber was assembled on a 22 × 40 mm^2^ glass cover slide with a pair of parallel copper strips (3M, Cergy-Pontoise, France) located 1 mm apart from each other (see Figure S6b). Small Parafilm pieces were attached onto the glass slide 10 mm apart from each other to seal the ends and form a closed compartment. An aliquot of 50 µL GUV solution 1:1 diluted in 105 mM glucose buffer was added in the spacing between the copper tapes and a 22 × 22 mm^2^ cover slide was placed on top of the solution.

Contrary to the Eppendorf chamber, the thickness of the home-made chamber is comparable to the one used for the rest of microscopy experiments. However, the home-made chamber could not be used together with the external UV LED on the confocal microscope due to safety reasons associated with strong reflection of the UV light from the copper tapes to the user. By repeating the electrodeformation experiments with both chambers under a phase contrast microscope, which does not require the attachment of external UV LED, we could rule out the effect of chamber thickness on the penetration of UV light into the sample (Figure S8c).

The copper tape electrodes were connected to Agilent 33220A 20 MHz Function/Arbitrary Waveform Generator (Agilent Technologies, USA) as shown in Figure S6. The voltage and frequency of the electric field were set to 10 V (peak to peak) and 1 MHz, respectively.

Before the electric field application, a typical, tensionless GUV adopts a quasi-spherical morphology and displays visual membrane fluctuations through the microscopy observations. In the vesicle electrodeformation method, a mild AC-field is used to pull out the excess vesicle area stored in membrane fluctuations and tubes by deforming the vesicles into ellipsoidal shapes [29] thus providing an accurate and direct assessment of total membrane area. The area of an ellipsoidal vesicle is:

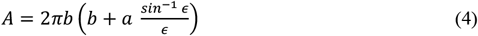

Here, *a* and *b* are the vesicle semi-axes along and perpendicular to the applied electric field, while *ϵ* denotes to ellipticity: *ϵ*^2^ = 1 − (*b*/*a*)^2^.

After recording the area increase of GUVs under AC-field, they were next illuminated with UV light while the AC-field was also still switched on and further area increase of the vesicles were recorded for 40-50 seconds at an acquisition speed of 8 frames per second (fps), see Figure S7. Subtracting the initial vesicle area in the absence of UV light (but with electric field on), *A*_*i*_, from the vesicle area under UV light, *A*_*UV*_, yields the percentage of relative area increase as 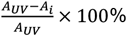 related to the *trans-*to*-cis* photoisomerization of azo-PC.

The length of the vesicle semi-axes was measured from the recorded vesicle images using Fiji software. Between 10 and 15 GUVs were analyzed from 3 separate sets of experiments for each condition to generate statistics. The corresponding plots in Figure S7 and Figure S8 are prepared with Origin Pro. The statistical significance of the vesicle area changes from different microscopy techniques and chamber conditions was tested with the one-way analysis of variance (ANOVA) and T-test (*p*-values for null-hypothesis were found as 0.76 and 0.082, respectively).

Since the majority of observations in this manuscript relies on confocal microscopy imaging performed in a thinner home-made chamber, electrodeformation calculations were also performed through confocal images focused on the equatorial trajectories of the vesicles sampled on the home-made chamber with same thickness dimensions as in the rest of the experiments. Because confocal images display information from a single focal plane, any potential experimental error or deviation was checked carefully and the accuracy of the confocal experiments to obtain the precise maximum projection of the vesicles for area increase calculations was compared to the results from phase contrast microscopy (Figure S8c). Similarly, effects from difference in the chamber thickness potentially affecting the UV irradiation through the GUV sample were also examined (Figure S8c).

### Analysis of changes in the apparent light-induced and adhesion areas

In order to check the relation between the UV-induced area increase of GUVs and the increase in the membrane area adhered to the protein condensates, the interaction of GUVs and protein condensates were monitored first in the absence and then in the presence of UV light under confocal microscopy. Subtracting the initial vesicle area in the absence of UV irradiation from the vesicle area under UV illumination allowed us to deduce the apparent area increase of azo-PC GUVs interacting with the protein condensates. The observed area increase is associated with the *trans-*to*-cis* photoisomerization of the azo-PC molecules. We defined the area increase of the spherical segments as ‘apparent area increase’ and further compared it to the expected area increase as assessed from electrodeformation method.

Statistics of the plots in Figure 6 in the main text and Figure S9 were generated with 11 data points for ten pairs of GUV-condensate systems in which each vesicle was interacting with only one condensate. To calculate the areas of the bare membrane segment and the one in contact with the condensate, we assumed spherical cap geometry. All plots were generated through Origin Pro software.

In order to check the correlation of the size differences between the interacting GUV and condensate to the distribution of the adhered area changes in the above-mentioned plots, we analyzed the quadratic proportionality of the radius of the interacting condensate to the radius of the GUV as 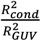 and the resulting values are displayed above each datapoint at Figure S9. Based on the indicated results, no correlation was detected between these two parameters and the distribution of datapoints among the plot in Figure S9.

## Supporting Information

Supporting Information is available from the Wiley Online Library or from the authors.

## Authors Contributions

A. M. and M. A. contributed equally to this work. R. D. designed the project. A.M., M.A. and M.S. performed the experiments and analyzed the data. R. L. and R.D. developed the theoretical interpretation. All authors wrote and edited the manuscript.

## Acknowledgements

A. M. acknowledges support from the Alexander von Humboldt foundation. M. A. acknowledges funding from the International Max Planck Research School on Multiscale Bio-systems. The authors would like to acknowledge Dr. Nannan Chen for providing the purified protein.

