## Supporting information for "Photoswitchable endocytosis of biomolecular condensates in giant vesicles"

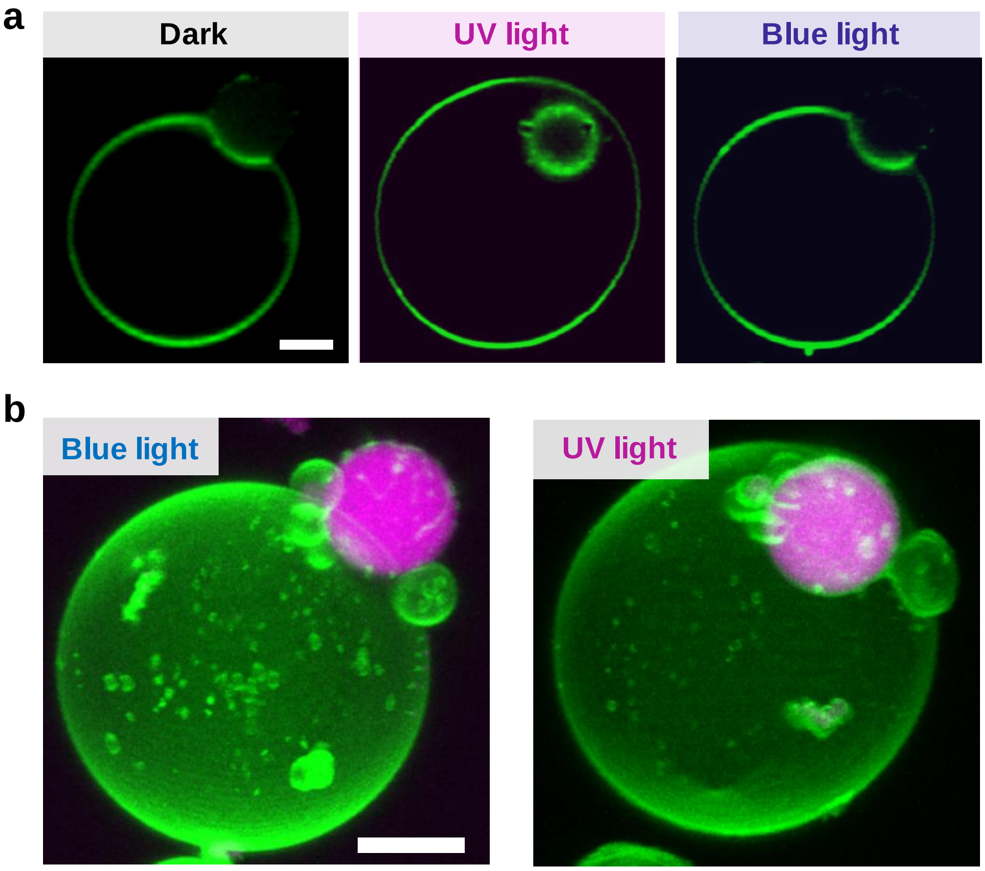


**Figure S1.** (a) Membrane channel and (b) 3D confocal projection for the vesicle shown in Figure 2b under blue and UV light as indicated. Note that the centers of the condensate droplet and the vesicle are not always in focus. Thus, contrary to 3D projections, confocal cross-sections do not always show properly the area increase and degree of engulfment upon isomerization. Scale bars are 5 µm.

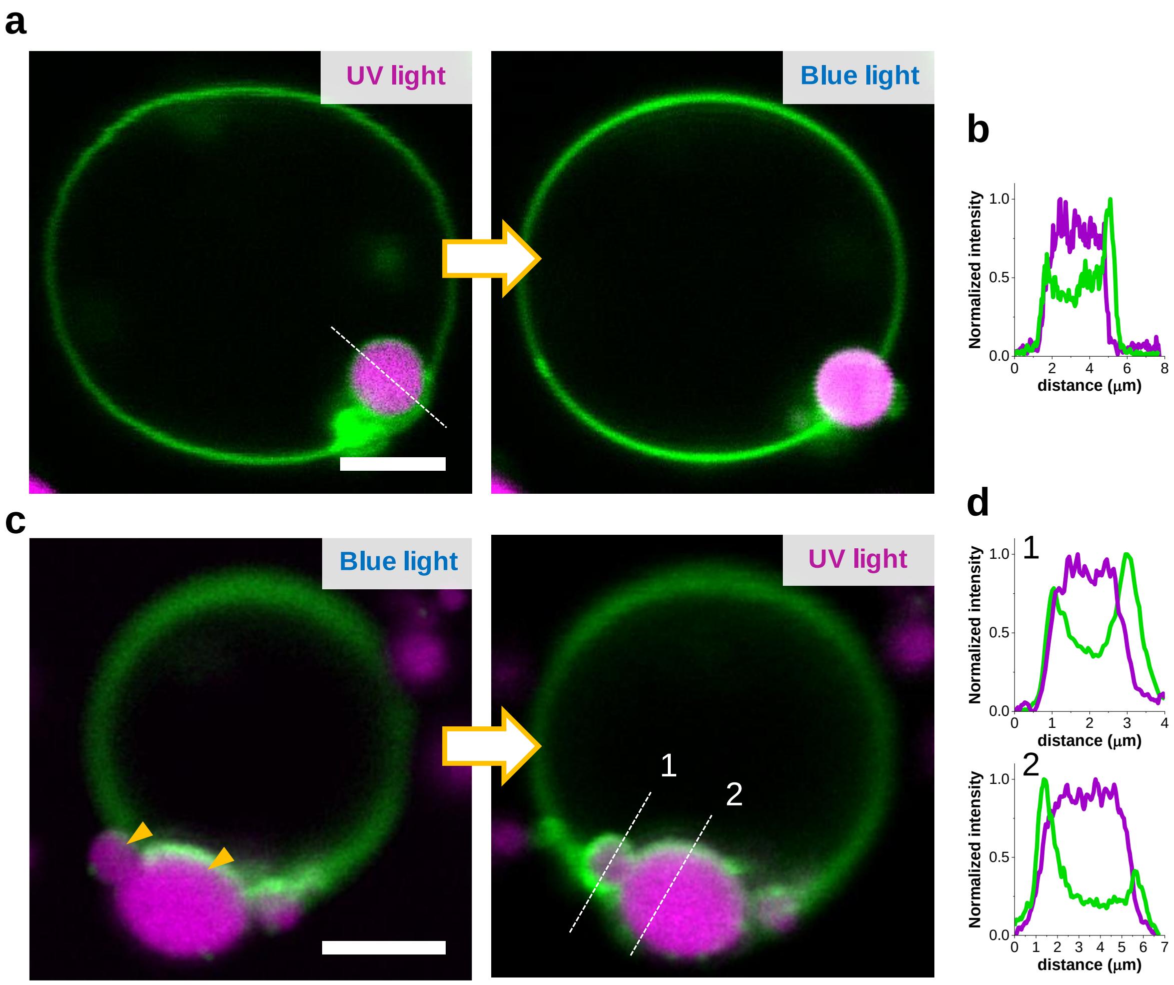


**Figure S2.** Confocal microscopy images showing examples of photoswitchable endocytosis. GUVs composed of equimolar POPC and azo-PC and labelled with 0.1 mol% Atto-647N-DOPE. Glycinin condensates are labelled with 10 µM SRB. (a) The completely engulfed condensate is released after blue light exposure. (b) Intensity profiles (green – membrane, magenta – condensate) for the dashed line shown in (a) demonstrate that the droplet is fully wrapped by the membrane evidenced by the two peaks in the green profile. (c) The condensates indicated by the yellow arrows get engulfed after UV light exposure. (d) Intensity profiles for the dashed lines shown in (c). Scale bars are 5 µm. Further examples for complete engulfment are found in Fig S4.

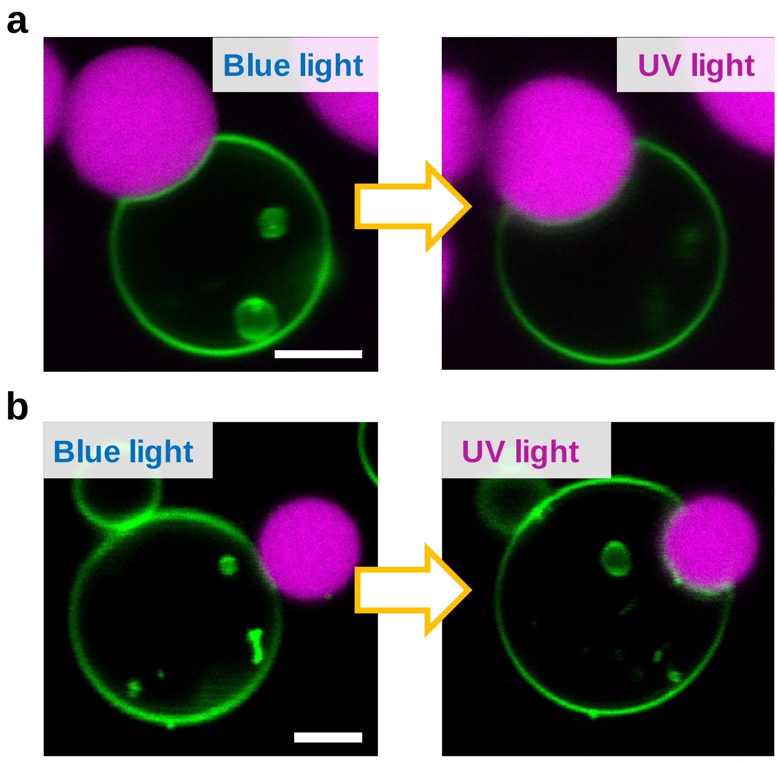


**Figure S3.** (a-b) Confocal microscopy images showing examples of photoswitchable partial-engulfment. GUVs composed of equimolar POPC and azo-PC and labelled with 0.1 mol% Atto-647N-DOPE. Glycinin condensates are labelled with 10 µM SRB. Scale bars are 5 µm.

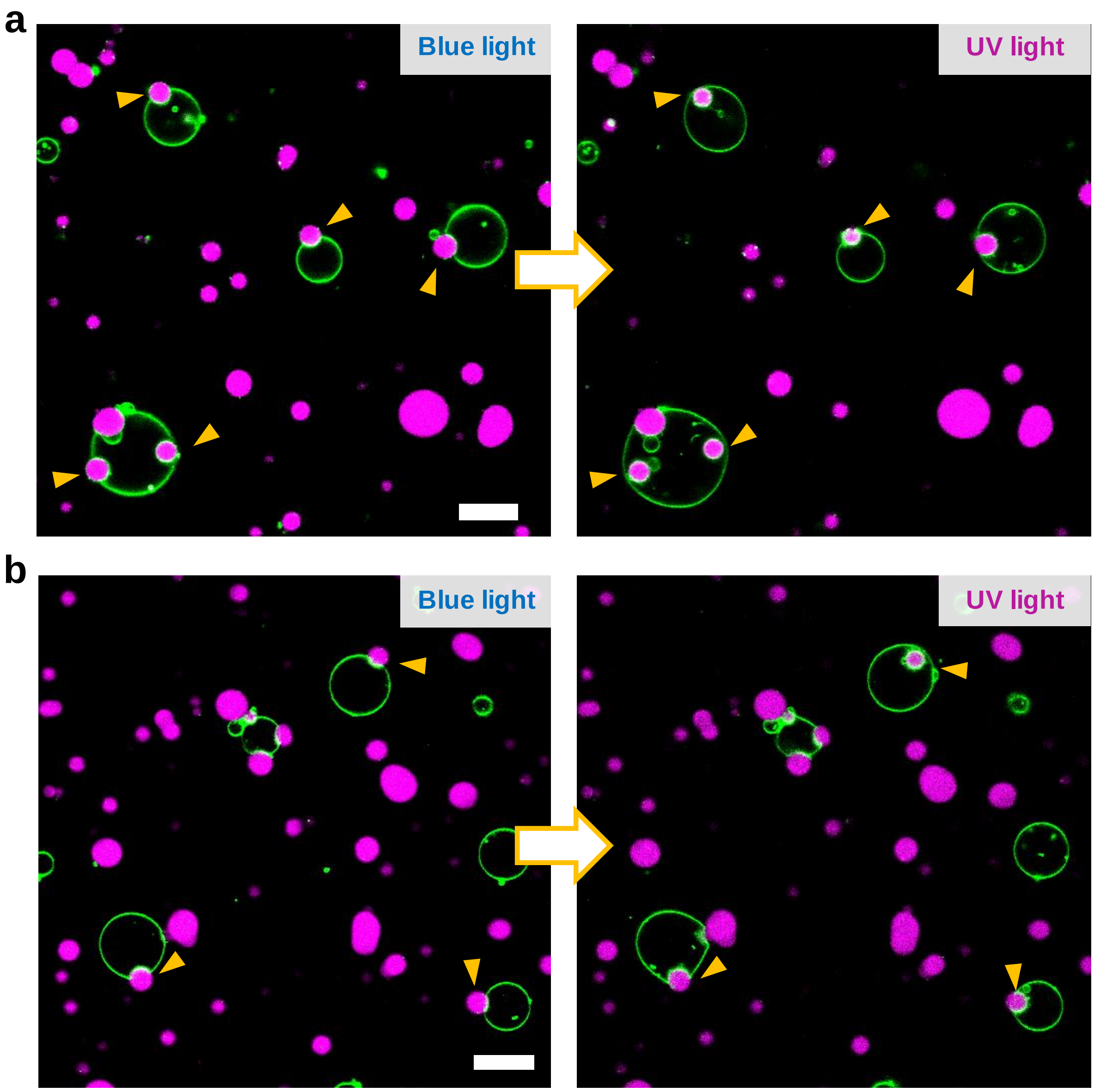


**Figure S4.** Confocal fluorescence full field images taken under UV/blue light exposure as indicated. The GUVs are composed of equimolar POPC and azo-PC and labelled with 0.1 mol% Atto-647N-DOPE. The glycinin condensates are labelled with 10 µM SRB. The yellow arrowheads point to the reversibly engulfed condensates, whether partially or fully. Scale bars are 20 µm.

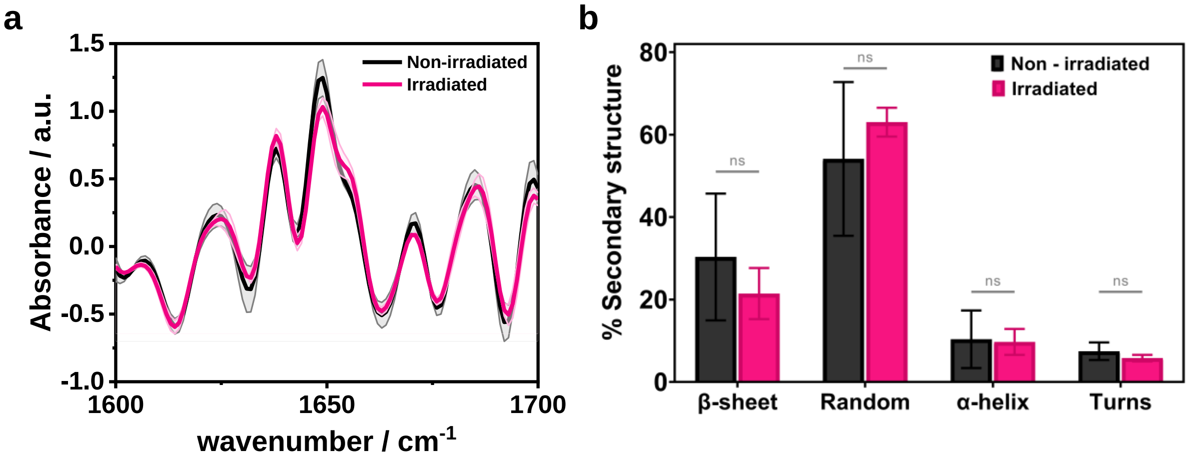


**Figure S5.** FTIR-ATR analysis of the secondary structure content of glycinin condensates before and after irradiation with UV-light. **a.** Second derivative of glycinin condensates at 150 mM NaCl before and after irradiation with UV-light for 60 seconds (note that much shorter irradiation times were used in the vesicle-condensate studies). **b.** Percent of secondary structure change before and after irradiation with UV-light. There are no significant differences between the irradiated and non-irradiated samples.

**Table S1.** Full width at half maximum (FWHM) input values in $cm^{-1}$ and their physically plausible ranges expected for each type of secondary structure as measured with ATR-FTIR.[1]

| **Secondary structure component** | | **FWHM input ($cm^{-1}$)** | **Lower limit ($cm^{-1}$)** | **Upper limit ($cm^{-1}$)** |
|---|---|---|---|---|
| *β-sheet* | *High wavenumber component (1670 – 1680)* | 9 | 8 | 11 |
| | *Low wavenumber component (1610 – 1640)* | 22 | 11 | 33 |
| *Random (1640 – 1650)* | | 55 | 50 | 60 |
| *α-helix (1650 – 1660)* | | 20 | 5 | 30 |
| *Turns (1660 – 1670; 1680 – 1700)* | | 20 | 5 | 30 |

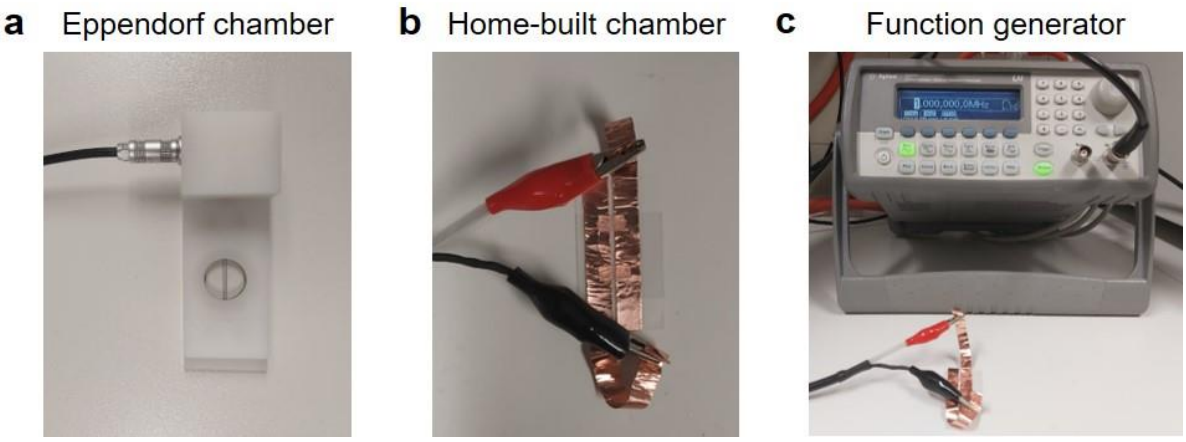


**Figure S6:** Pictures of electrodeformation chambers before being mounted on a microscope: (a) Eppendorf electrofusion chamber and (b) home-built chamber for electrodeformation assembled from two coverslips sandwiching copper-tape electrodes and parafilm strips. (c) Function generator to apply AC field to induce electrodeformation of GUVs. The two different chambers were used to compare the effects of sample thickness because of concerns regarding the penetration of the UV light.

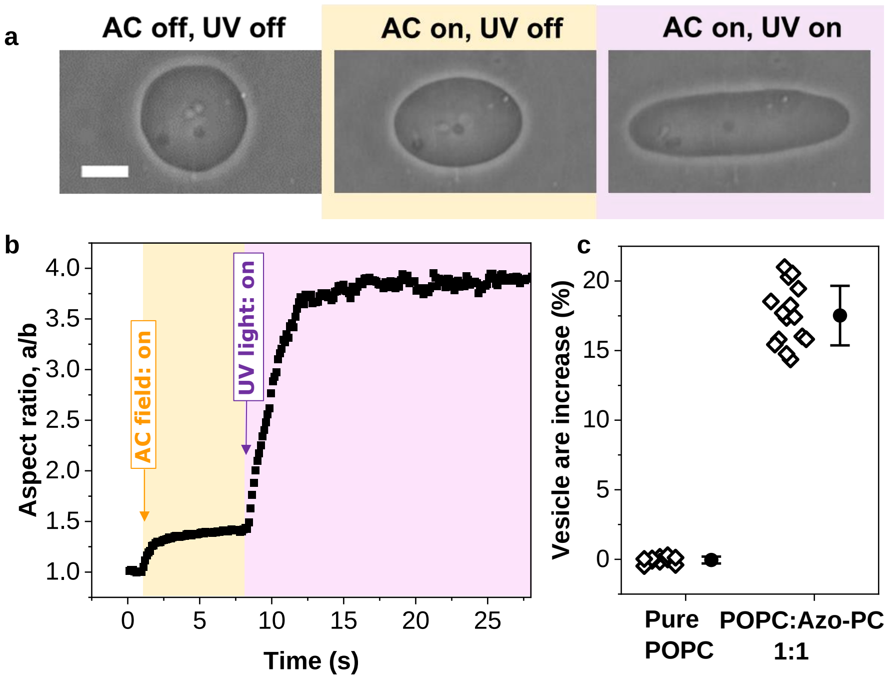


**Figure S7:** (a) Phase contrast images of a GUV composed of POPC:azo-PC (1:1) in AC field and UV irradiation, as indicated. The arrow indicated the direction of the field. The vesicle was first exposed to an AC field (5 kV $m^{-1}$ and 1 MHz) to pull out thermal fluctuations and deform them into a prolate ellipsoid with semi-axes a and b. Then, while keeping the AC field on, UV irradiation (365 nm) was initiated. Scale bar is 10 μm. (b) Electrodeformation analysis of the GUV in (a). (c) Quantification of the UV-induced area increase of POPC and POPC:azo-PC (1:1) GUVs via vesicle electrodeformation. 10-15 GUVs per condition were analyzed from 3 separate sets of experiments. Each open diamond represents the result of single GUV analysis while the solid circles and line bars are mean values and standard errors. Azo-PC containing GUVs showed ~18±2% of area increase under UV illumination.

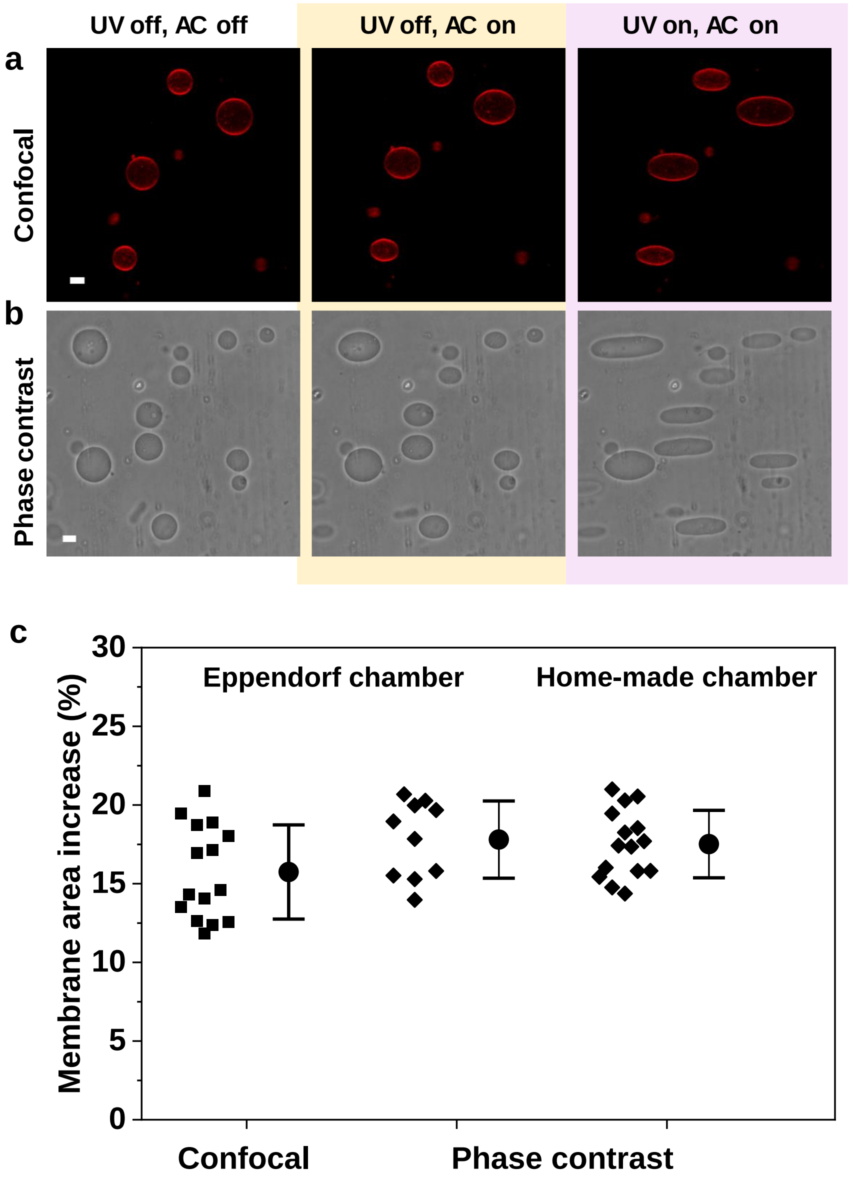


**Figure S8.** (a) Confocal and (b) phase contrast images of 50 mol% azo-PC containing GUVs during electrodeformation analysis. Initial state of GUVs before the exposure to AC field and UV light is demonstrated on the left panels. The middle panels show the morphology of GUVs in AC field only. In the right panel, GUVs are exposed to UV illumination as well as AC field. In order to monitor vesicles in confocal microscopy, GUVs are labelled with 0.1 mol% Atto-647N-DOPE. The scale bars are 10 µm. Quantification of light-induced membrane area under confocal and phase contrast microscopy based on vesicle electrodeformation. (c) The deformation of the vesicles in AC-field and UV light are measured through the changes in the vesicle aspect ratio. Membrane area of the GUVs are calculated through the area of the ellipsoid. By subtracting the initial membrane area in AC field from the membrane area under the influence of both UV light and AC field, we could obtain the UV induced area increase in the membrane. In addition to different microscopes, the effects of usage of Eppendorf and home-made chambers are also compared. Based on the ANOVA and T-tests ($p$-values are 0.76 and 0.082, respectively), the differences between different methods are not significant. Each filled square and diamond symbol represents the analysis of a single GUV while filled circles and line bars are the mean values and standard deviation. The average area increase of vesicles under UV illumination shows no dependence on the microscope mode or the used experimental chamber.

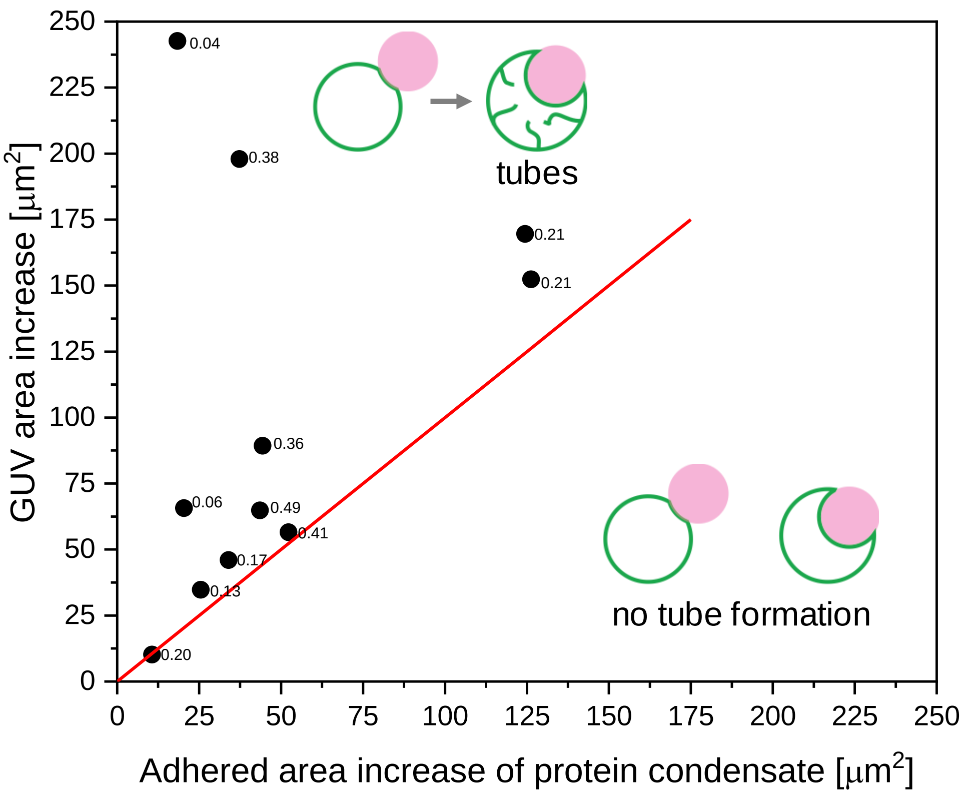


**Figure S9.** UV-induced GUV area increase versus adhered area increase of the glycinin condensates. GUVs composed of 1:1 POPC:Azo-PC and labelled with 0.1 mol% Atto-647N-DOPE. The adhered area changes detected through confocal screenshots in the absence and presence of UV irradiation were measured through FiJi software. The calculations were based on the spherical cap geometries. Each filled square is an individual datapoint generated from the pair of interacting GUV-condensate system in which a single vesicle interacted with a single condensate. The quadratic proportionality of the radius of the interacting condensate to the radius of the GUV, $\frac{R_{cond}^2}{R_{GUV}^2}$, were indicated on top of the each datapoint. Red line corresponds to the y=x line representing the datapoints when all the excess area accumulates on the membrane-condensate interface. The datapoints are distributed above the y=x line and their distribution does not show any correlated, common trend with the changes in $\frac{R_{cond}^2}{R_{GUV}^2}$, thus revealing that the adhesion changes are not correlated to size differences of the interacting GUV-protein condensate pairs and not all the UV-induced excess area of GUVs completely accumulated on the GUV-condensate interface to trigger to the adhesion process of the glycinin condensates.

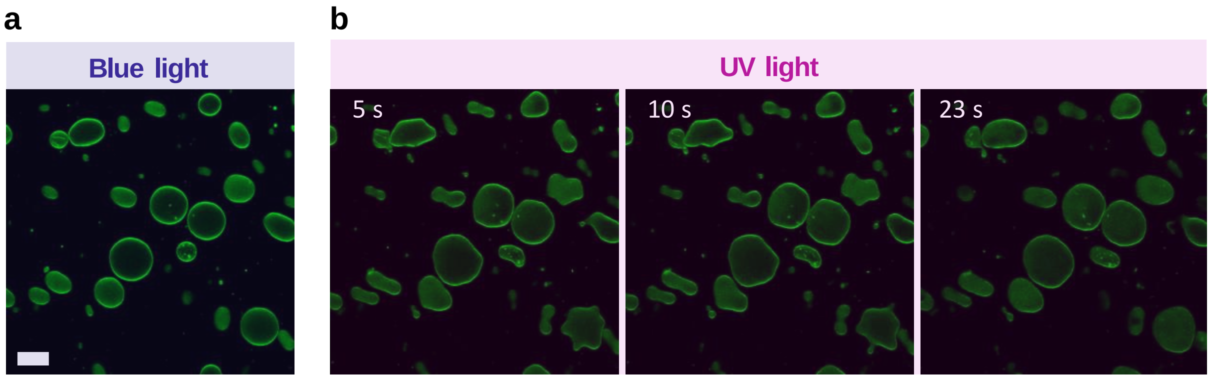


**Figure S10.** Blue (a) and UV (b) light induced shape transformations of 50 mol% azo-PC containing GUVs in low concentrations of sugar solution and in the absence of any salt asymmetry between internal and external GUV media. GUVs labeled with 0.1 mol% Atto-647N-DOPE were grown in 100 mM sucrose solution and 1:1 diluted to 105 mM glucose solution in absence of any salts. Upon UV irradiation (365 nm), the GUVs underwent complex shape transformations over time which were distinct from the internal tubulation events observed in the presence of high salt asymmetry like in Figure 1c in the main text. The time stamps of changing UV-induced morphological transitions of GUVs are shown in the upper part of the images. The scale bar is same for all these confocal images and corresponds to 10 µm.

### Movie Captions

**Movie S1:** GUVs composed of POPC:Azo-PC 1:1 and labelled with 0.1 mol% Atto-647N-DOPE were sequentially exposed to UV and blue light. The time stamps show the time in seconds after initiating acquisition. The scale bar is 50 µm.

**Movie S2:** Photo-switchable partial engulfment of a condensate under UV and blue light as indicated on the upper part of the video. The GUV is composed of POPC:Azo-PC 1:1 and labelled with 0.1 mol% Atto-647N-DOPE and the glycinin condensate was labelled with 10 µM SRB. The time stamps show the time in seconds after initiating acquisition. The scale bar is 5 µm.

**Movie S3:** Photo-switchable endocytosis of a condensate. The GUV composed of POPC:Azo-PC 1:1 and labelled with 0.1 mol% Atto-647N-DOPE in contact with the glycinin condensate (labelled with 10 10 µM SRB) was sequentially exposed to UV and blue light as indicated on the upper part of the video. The time stamps show the time in seconds after initiating acquisition. The scale bar is 5 µm.

**Movie S4:** Large field confocal image showing several examples of complete and partial condensate engulfment. The GUVs composed of POPC:Azo-PC 1:1 and labelled with 0.1 mol% Atto-647N-DOPE in contact with the glycinin condensate (labelled with 10 10 µM SRB) were sequentially exposed to UV and blue light as indicated in the corresponding frame of the video. The time stamps show the time in seconds after initiating acquisition. The scale bar is 20 µm.

**Movie S5:** Release of a completely engulfed condensate in blue light. The GUV composed of POPC:Azo-PC 1:1 and labelled with 0.1 mol% Atto-647N-DOPE were sequentially exposed to UV and blue light. Glycinin condensate was labelled with 10 µM SRB. The time stamps show the time in seconds after initiating acquisition. The periods of irradiation of the sample are indicated on the upper part of the video. The scale bar is 5 µm.

**Movie S6:** Engulfment of multiple condensates in UV light. The GUV composed of POPC:Azo-PC 1:1 and labelled with 0.1 mol% Atto-647N-DOPE was exposed to UV light. Glycinin condensates were labelled with 10 µM SRB. The time stamps show the time in seconds after initiating acquisition. The periods of irradiation of the sample are indicated on the upper part of the video. The scale bar is 5 µm. The video was processed with LAS X and Fiji software.

**Movie S7:** Condensate partial engulfment when exposed to UV light, displaying a large change in the contact angle between the condensate and the membrane. The GUV composed of POPC:Azo-PC 1:1 and labelled with 0.1 mol% Atto-647N-DOPE was exposed to UV light. Glycinin condensate was labelled with 10 µM SRB. The time stamps show the time in seconds after initiating acquisition. The periods of irradiation of the sample are indicated on the upper part of the video. The scale bar is 5 µm.

**Movie S8:** Condensate partial engulfment in UV light. The GUV composed of POPC:Azo-PC 1:1 and labelled with 0.1 mol% Atto-647N-DOPE was exposed to UV light. Glycinin condensate was labelled with 10 µM SRB. The time stamps show the time in seconds after initiating acquisition. The periods of irradiation of the sample are indicated on the upper part of the video. The scale bar is 5 µm.

[1] a) E. Goormaghtigh, V. Cabiaux, J.-M. Ruysschaert, *European Journal of Biochemistry* **1990**, *193* (2), 409, https://doi.org/https://doi.org/10.1111/j.1432-1033.1990.tb19354.x; b) G. Long, Y. Ji, H. Pan, Z. Sun, Y. Li, G. Qin, *International Journal of Food Properties* **2015**, *18* (4), 763, https://doi.org/10.1080/10942912.2014.908206.
